# Platelet HMGB1 in Platelet-Rich Plasma Promotes Tendon Wound Healing

**DOI:** 10.1101/2021.04.22.440884

**Authors:** Jianying Zhang, Feng Li, Tyler Augi, Kelly M. Williamson, Kentaro Onishi, MaCalus V. Hogan, Matthew D. Neal, James H-C. Wang

**Affiliations:** MechanoBiology Laboratory, Department of Orthopaedic Surgery; Department of Bioengineering; Department of Physical Medicine and Rehabilitation; Department of Surgery, University of Pittsburgh, Pittsburgh, PA15213

**Keywords:** Tendon injury, Wound healing, PRP, Platelets, HMGB1

## Abstract

Platelet-rich plasma (PRP) is a widely used autologous treatment for tendon injuries in clinics, but clinical trials often produce conflicting results. Platelets (PLTs) are a major source of high mobility group box1 (HMGB1) that is gaining attention as a chemoattractant that can recruit stem cells to the wound area to enhance healing; however, the contribution of PLT HMGB1 in wounded tendon healing remains unexplored. This study investigated the effect of PLT HMGB1 within PRP to enhance healing in an acute patellar tendon injury model in PLT HMGB1 knockout (KO) mice and GFP mice. A window defect was created in the patellar tendons of both groups of mice, and wounds were treated with either saline, PRP isolated from PLT HMGB1 KO mice, or PRP isolated from GFP mice. Seven days post-treatment, animals were sacrificed and analyzed by gross inspection, histology, and immunostaining for characteristic signs of tendon healing and repair. Our results showed that in comparison to mice treated with PRP from PLT HMGB1-KO mice, wounds treated with PRP from GFP mice healed faster and exhibited a better organization in tendon structure. Mice treated with PRP from PLT HMGB1-KO mice produced tendon tissue with large premature wound areas and low cell densities. However, wounds of PLT HMGB1 KO mice showed better healing with PRP from HMGB1 KO mice compared to saline treatment. Moreover, wounds treated with PRP from GFP mice had increased extracellular HMGB1, decreased CD68, increased stem cell markers CD146 and CD73, and increased collagen III protein expression levels compared to those treated with PRP from PLT HMGB1 KO mice. Thus, PLT HMGB1 within PRP plays an important role in the healing of wounded tendon. Our findings also suggest that the efficacy of PRP treatment for tendon injuries in clinics may be affected by PLT HMGB1 within PRP preparations.

## Introduction

Tendon injuries to the Achilles and patellar tendons are prevalent in both occupational and athletics populations. Overall, injured tendon healing is slow and yields an inferior quality of tendon tissue that is prone to re-injury. Many therapeutic approaches including injection of autologous platelet-rich plasma (PRP) have been devised to manage tendon injuries (1, 2). The use of PRP is a popular option in the treatment of tendon injuries in orthopaedics and sports medicine (1, 2), however the efficacy of PRP treatment on tendon injuries, particularly in clinical trials, has been controversial. Several studies have reported that PRP can effectively treat tendon injuries (3-5), whereas others have shown the opposite with no improvement in pain or tendon function after PRP treatment (6-8). Thus, further investigation into the role of platelets (PLTs) in tendon wound healing is essential to understand the PLT action mechanism in PRP and improve the efficacy of PRP in treating tendon injuries.

Although lacking nuclei, platelets are a rich source of high mobility group box-1 (HMGB1), a highly conserved nuclear protein that is released by all cell types upon injury (9, 10). The function of HMGB1 as an inflammatory molecule or chemoattractant is dependent upon its redox state. When inside the cell, either in the nucleus or cytoplasm, HMGB1 is completely reduced as fully reduced HMGB1 (frHMGB1). Once released to the extracellular matrix, frHMGB1 is partially oxidized to disulfide HMGB1 (dsHMGB1) that is believed to initiate inflammation (11). Research has suggested that during platelet activation, HMGB1 is presented on the cell surface and released in significant amounts, with the current theory that HMGB1 is fully reduced at this stage (12-15). Therefore, we hypothesized that platelet HMGB1 (PLT HMGB1) within PRP may have an important role in tendon injury healing. To test the hypothesis, we investigated the effect of PLT HMGB1 within PRP on wounded tendon healing using a transgenic mouse line with platelet-specific ablation of HMGB1 (PLT HMGB1-KO). The findings of this study showed that PLT-HMGB1 within PRP preparations is able to facilitate proper healing and repair of tendon injuries.

## Materials and methods

### Animals

All experiments were performed according to the relevant guidelines and regulations. The protocol for animal use was approved by the Institutional Animal Care and Use Committee of the University of Pittsburgh (IACUC protocol #18083391). C57BL/6-Tg (UBC-GFP) mice were obtained from Jackson Laboratory (Bar Harbor, ME). Mice with platelet-specific ablation of HMGB1 (Pf4-Cre Hmgb1^fl/fl^ mice, referred to as PLT HMGB1-KO mice) were generated as described elsewhere using the Cre/loxP system (16).

### Isolation and preparation of platelets and PRP

Mice were anesthetized with isoflurane, and blood was drawn from the retro-orbital plexus into anti-coagulated tubes. PRP was obtained by centrifugation at 500g for 10 min. Platelets were first pelleted from PRP by centrifugation at 1,000g for 10 min, and were then resuspended in ACD buffer consisting of 39 mM citric acid, 75 mM sodium citrate, and 135 mM dextrose with 5 mM of EDTA according to the published protocol (17). This platelet solution was used for the following experiments.

### Determination of HMGB1 in activated platelets

Isolated platelets were activated by adding 100 μl of 10,000 U/ml bovine thrombin solution into 0.4 ml of 1 × 10^8^/ml platelet-ACD solution at room temperature for 30 min. The reaction mixture was centrifuged at 1,000g for 10 min, and the supernatant was collected to determine the amount of HMGB1 released from platelets using an HMGB1 ELISA kit according to the manufacturer’s protocol (Shino-Test Corporation, Tokyo, Japan). The pellet was re-suspended with 0.9% of sodium chloride solution and reacted with a rabbit anti-mouse HMGB1 primary antibody for 3 hrs at room temperature (1:350, abcam, Cat. #ab18256), followed by a goat anti-rabbit secondary antibody conjugated with Cy3 for 1hr at room temperature (1:500, Millipore, Cat. #AP132C).

### Mouse tendon wound healing model

The effect of PLT HMGB1 on tendon wound healing was tested with a window defect created in each patellar tendon (PT) of PLT HMGB1-KO mice and GFP mice using a 1 mm diameter biopsy punch. Wounded PLT HMGB1-KO mice and GFP mice were divided into three groups (3 mice/group). Group 1 mice were treated with 10 μl of saline (**Saline**), group 2 mice were treated with 8 μl of PLT HMGB1-KO PRP and 2 μl of bovine thrombin for PRP activation (**KO-PRP**), and group 3 mice were treated with lox 8 μl of GFP-PRP and 2 μl of bovine thrombin (**GFP-PRP**). All mice were sacrificed at 7 days post-injury, and patellar tendons were harvested. The effect of PLT HMGB1 on wounded tendon healing and cell migration was assessed by histological analysis.

### Histochemical staining on mouse tendon tissue sections

Tendon tissue sections were fixed with 4% paraformaldehyde for 20 min at room temperature, and then washed three times with PBS. Slides were stained with H&E at room temperature according to the standard protocols, washed with water 3 times, and dehydrated through 15%, 30%, 50%, 75%, 95% alcohol, and absolute alcohol for five minutes each. Finally, slides were treated with xylene and mounted with resinous mounting medium. The staining results were observed and imaged on a microscope (Nikon eclipse, TE2000-U).

### Immunostaining on mouse tendon tissue sections

For immunostaining, the patellar tendons were dissected from the mice and were immediately immersed in O.C.T compound (Sakura Finetek USA Inc., Torrance, CA) in disposable molds and frozen at -80°C. Then, cryo-sectioning was performed at -25°C to obtain ∼ 8 µm thick tissue sections, which were left at room temperature overnight. The tissue sections were fixed in 4% paraformaldehyde for 15 min and blocked with universal blocking solution (ThermoFisher Scientific, Pittsburgh, PA, Cat. #37515). The sections were then incubated with rabbit anti-mouse HMGB1 antibody (1:350, abcam, Cat. #ab18256) at 4°C overnight followed by goat anti-rabbit secondary antibody conjugated with Cy3 for 1 hr at room temperature (1:500, Millipore, Cat. #AP132C). Since the purpose of this staining was to evaluate the presence of extracellular HMGB1 in the tendon, the tissue sections were not treated with the penetration reagent Triton-X100 that permeates the nuclear membrane.

Similarly, the fixed tissue sections were reacted individually overnight at 4°C with the following primary antibodies: rabbit anti-CD68 antibody (1:500, Abcam, Cat. #125212, Cambridge, MA), rabbit anti-CD146 antibody (1:500, Abcam, Cat. #75769; Cambridge, MA), rabbit anti-CD73 antibody (1:500, LSBio, Cat. #LS-B14527-50, Seattle, WA), or rabbit anti-collagen III antibody (1:500, ThermoFisher, Cat. #22734-1-AP, Waltham, MA). In the next morning, tissue sections were washed 3 times with PBS and incubated at room temperature for 2 hrs with Cy3-conjugated goat anti-rabbit IgG antibody (1:500, Millipore, Cat. #AP132C).

Total cell numbers were stained with 4.6-diamidino-2-phenylindole (DAPI). The stained tendon tissue sections were imaged using the fluorescent microscope (Nikon eclipse, TE2000-U).

### Semi-quantification of positively stained tissue sections

The percentage of HMGB1 expression in activated platelets from PLT HMGB1-KO and GFP mice was determined by semi-quantification. Platelets were stained for HMGB1 (as above) from three mice of each group, smeared onto a glass slide, and imaged using a fluorescent microscope (Nikon eclipse, TE2000-U). The percentages of HMGB1 expression in platelets were calculated by dividing the number of positively stained platelets with red fluorescence by the total number of platelets.

For semi-quantification of cell marker staining in tissue sections, stained tissue sections (3 sections/mouse) in each group (3 mice/group) were examined under a microscope and five random images were taken. Positively stained areas were manually identified by examining the images taken and processed using SPOT™ imaging software (Diagnostic Instruments, Inc., Sterling Heights, MI). The proportion of positive staining was calculated by dividing the total area viewed under the microscope by the positively stained area. These values were averaged to represent the percentage of positive staining in all the groups.

### Statistical analysis

All results were obtained from 6 tendons (three mice) from each group and presented as the mean ± SD. The statistical analyses were performed with an unpaired student *t*-test. When P < 0.05, the two groups in comparison were considered significantly different.

## Results

### Expression of HMGB1 is decreased in platelets of PLT HMGB1-KO mice

First, the expression of PLT HMGB1 was assessed for both transgenic lines, specifically Pf4-Cre Hmgb1^fl/fl^ mice, referred to as PLT HMGB1-KO mice, and GFP mice. Platelets were isolated and stained as described in the methods. Immunostaining showed that PLT HMGB1 was decreased within PLT HMGB1-KO mice (**Fig. 1A, B**) compared to GFP mice (**Fig. 1C, D**). Semi-quantification (**Fig. 1E**) confirmed these results showing only ∼7% of platelets in PLT HMGB1-KO mice stained positively for HMGB1, but 86% of platelets in GFP mice stained positively for HMGB1. After verifying both transgenic lines for the level of PLT HMGB1, further experiments were performed using PRP preparations to investigate the role of PLT HMGB1 in healing and repair within PRP preparations.

**Fig. 1.**
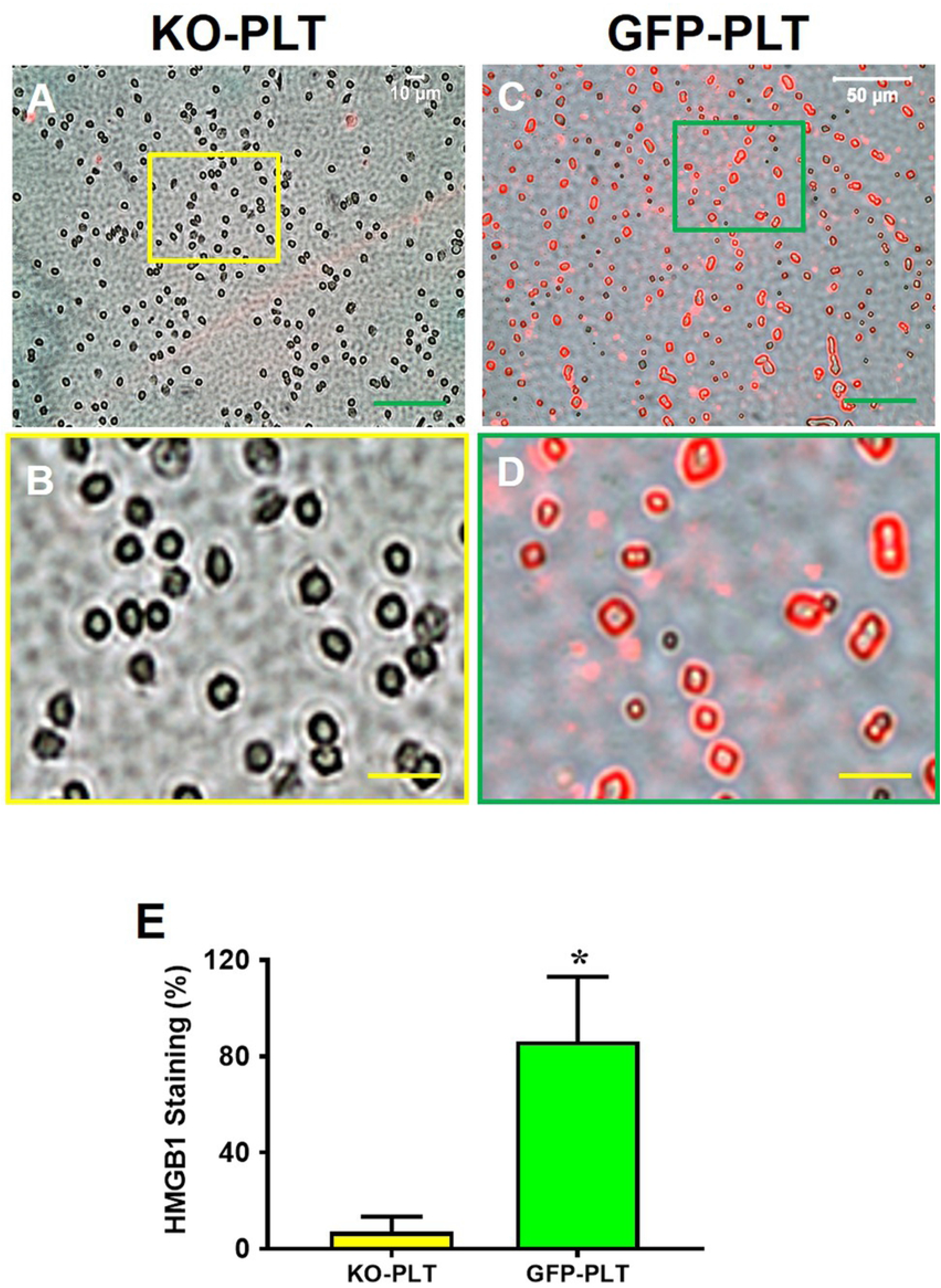
KO-PLTs have much less HMGB1 than GFP-PLTs. **A, B:** HMGB1 expression in the platelets of HMGB1-KO mice (KO-PLT); **C, D:** HMGB1 expression in the platelets of GFP mice (GFP-PLT); **B, D:** Enlarged images of the box areas in the image **A** and **C. E:** Semi-quantification of positively stained platelets with HMGB1 confirms the results of immunostaining showing that only a few (∼7%) platelets in HMGB1-KO mice are positively stained with HMGB1 compared to more than 86.8% of platelets in GFP mice were positively stained with HMGB1. *P < 0.01 (GFP-PLT vs. KO-PLT). Green bars: 50 μm; Yellow bars: 10 μm.

### PRP generated from PLT-HMGB1 KO mice impairs tendon wound healing

Patellar tendon (PT) wounds of mice from each group were treated individually with PRP generated from either PLT HMGB1-KO mice or from GFP mice. A saline treatment was used as a control treatment. Saline treated wound exhibited large unhealed wound areas (red arrows in **Fig. 2A, D**), whereas wounds treated with either KO or GFP-PRP in both mice groups (**Fig. 2B, C, E, F**) had better healing compared to saline treated wounds. Further gross inspection showed that wound healing in the patellar tendons of PLT HMGB1-KO mice displayed unhealed wound areas in all treatment groups (**Fig. 2A-C**, yellow and green arrow in **B** and **C**, respectively) compared to GFP mice (**Fig. 2E, F**). PLT HMGB1-KO mice treated with GFP-PRP (**Fig. 2C**) exhibited slightly better healing compared to treatment with KO-PRP (**Fig. 2B**). However, PLT HMGB1-KO mice treated with KO-PRP (**Fig. 2B**) had better healing than a saline treatment (**Fig. 2A**, red arrow). These results indicated that PLT HMGB1 within PRP preparations is required for promoting tendon wound healing by PRP.

**Fig. 2.**
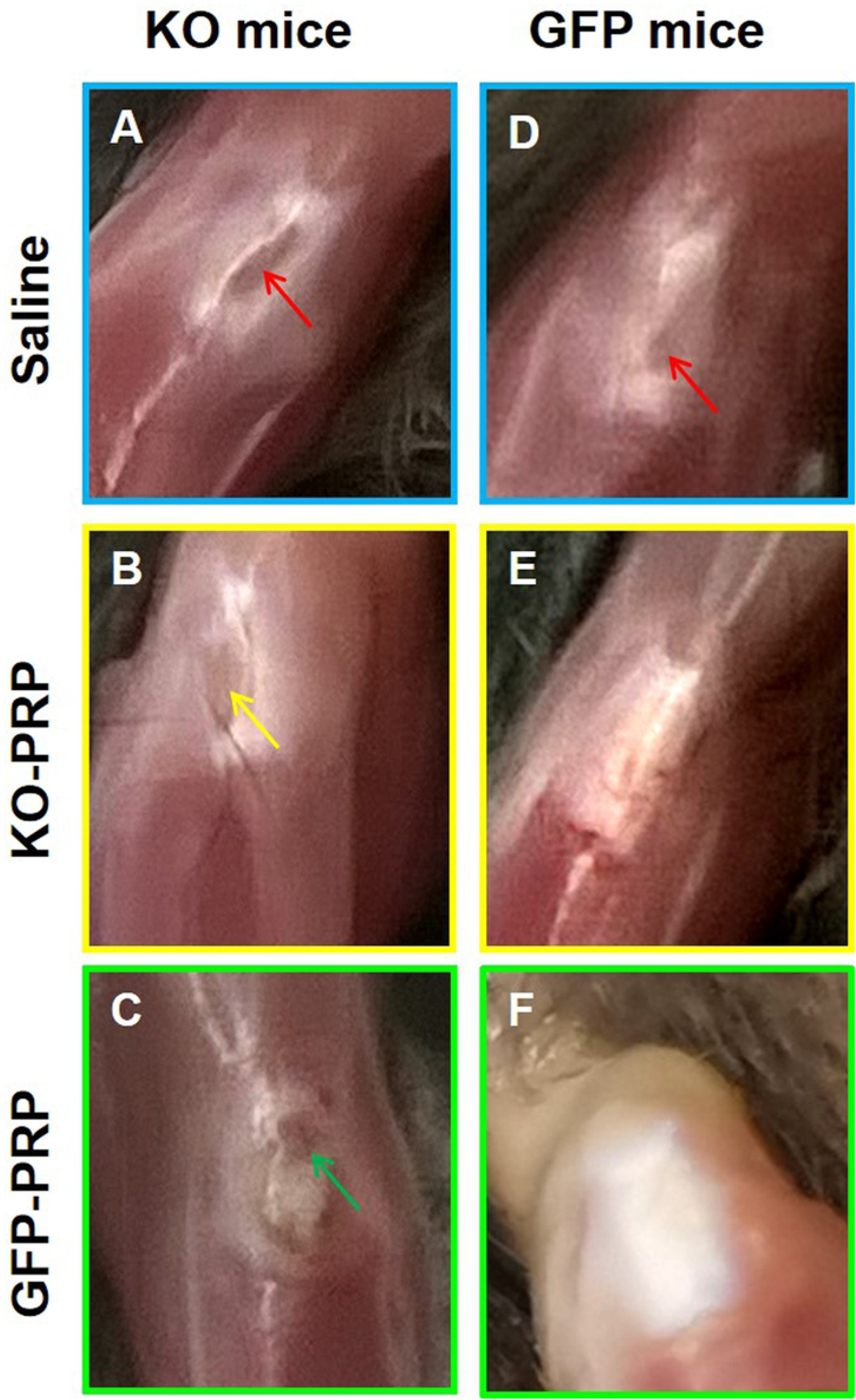
PRP generated from PLT HMGB1-KO mice adversely affects wounded PT healing. Results show that wounded PTs of PLT HMGB1-KO mice (KO mice) healed slower (**A-C**, red, yellow, and green arrows point to unhealed area) than PTs in GFP mice (**D-F**). However, PRP (KO-PRP or GFP-PRP) treated wound (**B, C, E, F**) healed much faster than the wounds treated with saline (**A, D**, red arrows). PT: patellar tendon; KO-PRP: PRP prepared from PLT HMGB1-KO mice; and GFP-PRP: PRP prepared from GFP mice.

Histological analysis by H&E staining confirmed our gross inspection results. The wounds in the patellar tendons of PLT HMGB1-KO mice healed poorly overall, while wounds in GFP-PRP treated mice healed faster (**Fig. 3**). Specifically, PLT HMGB1-KO mice treated with KO-PRP still exhibited large unhealed areas (black arrows, **Fig. 3B**) although GFP-PRP treated KO mouse wound displayed better healing (**Fig. 3C**) compared to that treated with KO-PRP (**Fig. 3B**). In contrast, GFP-mice treated with GFP-PRP had complete healing with a normal-like tendon appearance (**Fig. 3F**), while unhealed wound areas were found in GFP mice treated with KO-PRP (**Fig. 3E**). GFP-PRP treated wounds in the patellar tendon of GFP mice displayed newly formed tendon tissue in the wound area with normal-like tendon organization (**Fig. 3F**). Saline treated tendons produced large unhealed wound areas in both mice groups (red arrows in **Fig. 3A, D**), in contrast to PRP treatment which generally enhanced healing in patellar tendons despite the model or form of PRP used as a treatment (**Fig. 3B-F**). These results further confirm that PLT HMGB1 within PRP preparations is able to enhance tendon healing, and without PLT HMGB1, tendon wound healing is reduced and slowed in comparison.

**Fig. 3.**
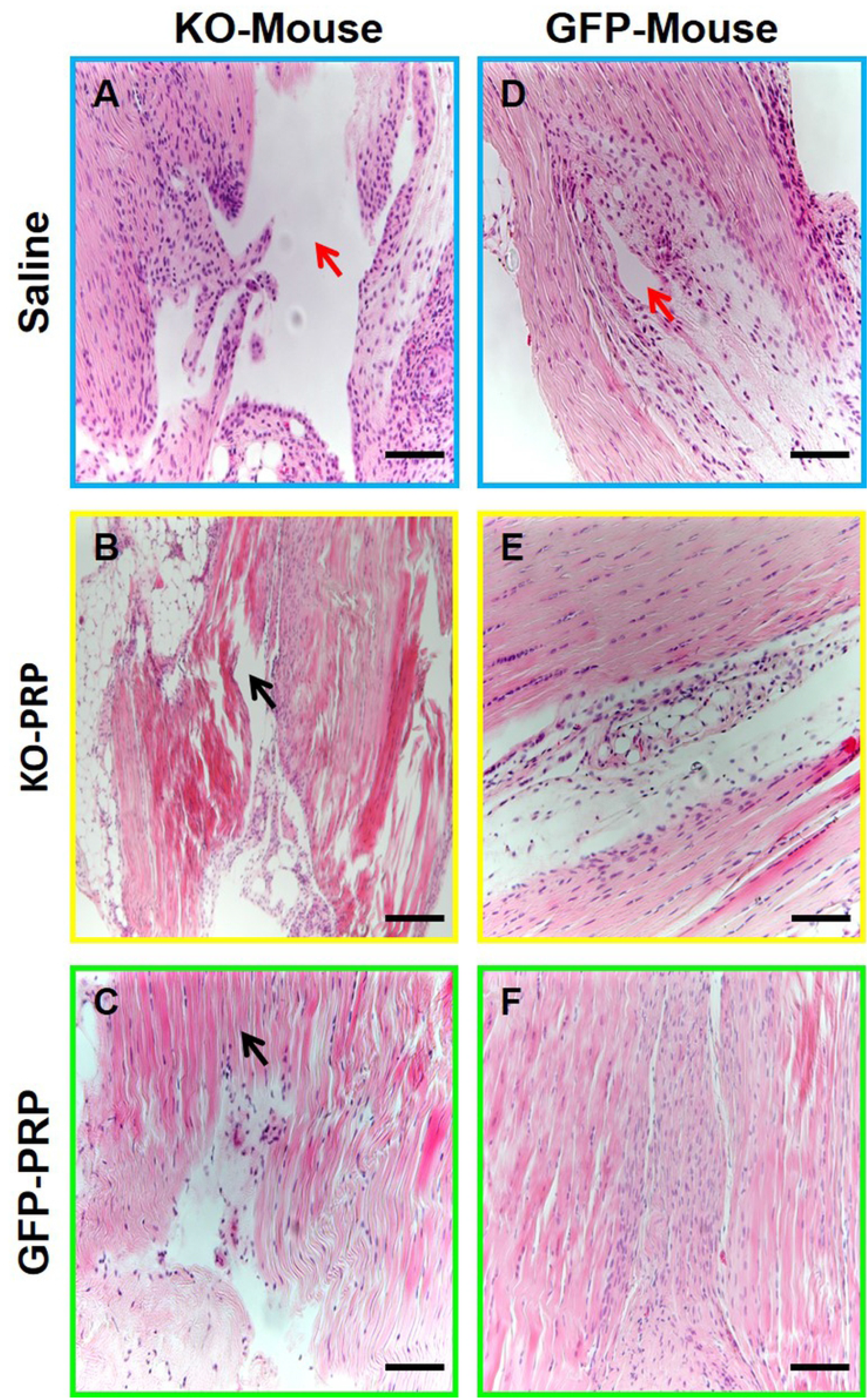
Wounded PTs treated with KO-PRP heals much slower than those treated with GFP-PRP. Results show that wounded patellar tendons in PLT HMGB1-KO mice heal slower (**A-C**, black arrows in **B** and **C** show unhealed and disorganized area) than GFP mice (**D, F**). PRP treated wounds (**B, C, E, F**) healed much faster than wounded patellar tendons treated with saline (**A, D**, red arrows point to large unhealed area). KO-PRP: PRP from PLT HMGB1-KO mice; and GFP-PRP: PRP from GFP mice. Black bars: 100 μm (H&E staining).

### HMGB1 level in tendon matrix is low in PLT-HMGB1-KO mice after PRP treatment

Previous research has shown that the release of local HMGB1 from injured tissues can enhance the healing and regeneration of skeletal, hematopoietic and muscle tissues *in vivo* (18). Thus, to further evaluate the role of PLT-HMGB1 in PRP preparations in healing and repair, the patellar tendons from each transgenic injury model with their respective PRP treatments were evaluated for the release of tendon tissue specific HMGB1 at the site of injury. Each patellar tendon was assessed with immunostaining for HMGB1 to determine how the presence or absence of PLT HMGB1 in both the GFP-PRP and KO-PRP may affect healing and repair within our model. Our results showed that the acute injury model induced the release of HMGB1 from local tendon tissue surrounding the wound area to the tendon matrix in both GFP mice and PLT HMGB1-KO mice, as evidenced by positively stained HMGB1 (red fluorescence in **Fig. 4**). However, reduced levels of HMGB1 can be seen within the tendon matrix of PLT HMGB1-KO mice (**Fig. 4A-L**) in comparison to the elevated level of HMGB1 released in tendons of GFP mice (**Fig. 4M-X**). Fluorescent image analysis indicated that GFP-PRP treatment increased the levels of HMGB1 in the tendon matrix of both transgenic mouse lines specifically due to treatment with GFP-PRP (**Fig. 4I-L, 4U-X**). However, the concentration of locally released HMGB1 in the tendons treated with PLT HMGB1-KO-PRP were not significantly increased (**Fig. 4E-H**). Thus, PLT HMGB1 is able to enhance the presence of local HMGB1 at the injury site.

**Fig. 4.**
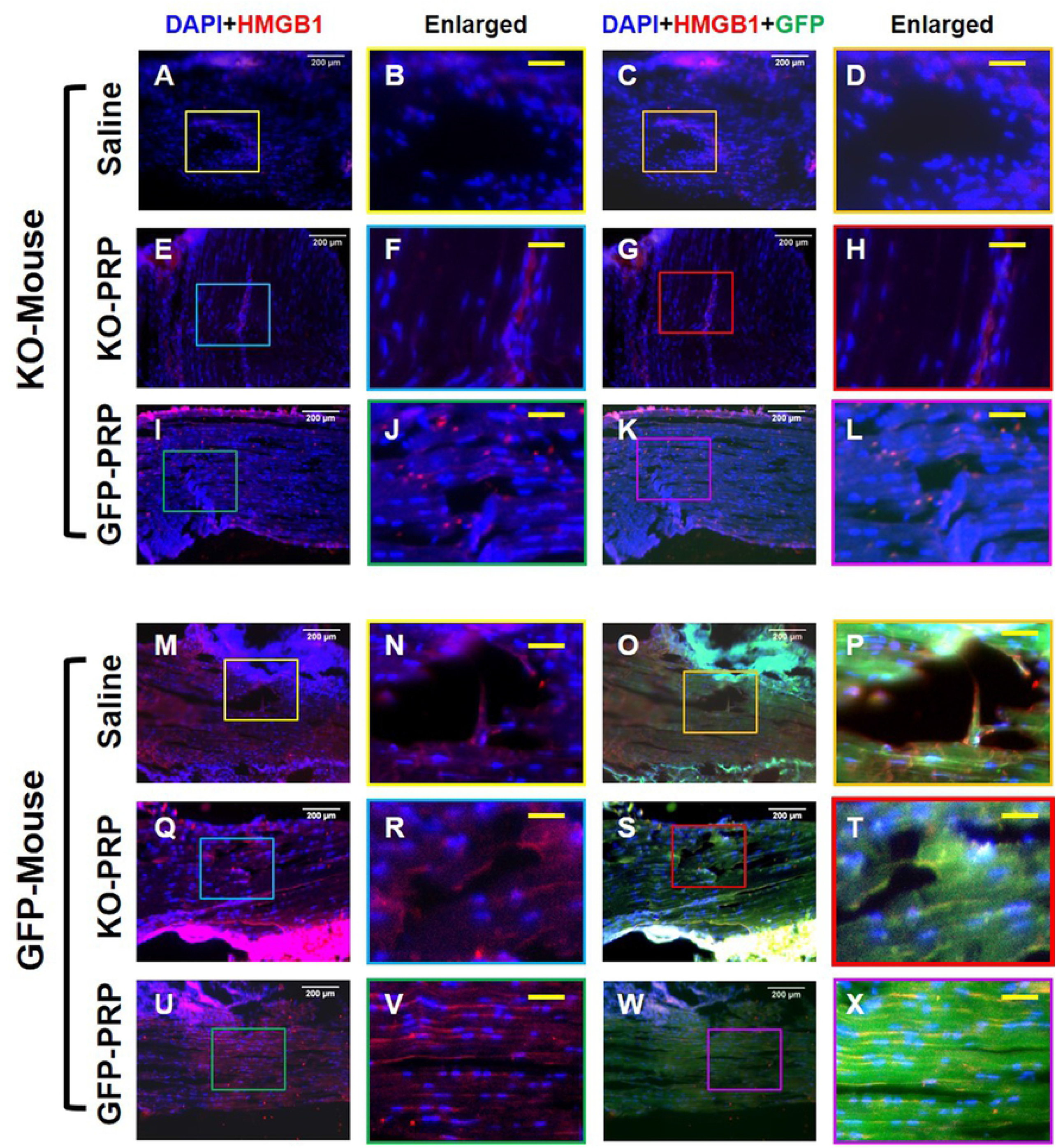
HMGB1 expression in KO-PRP treated wound is much lower than GFP-PRP treated wound. Higher levels of HMGB1 are found in the wound areas of GFP tendons (**M-X**) than that of PLT HMGB1-KO mouse patellar tendons (**A-L**). GFP-PRP treated wound areas (**I-L**, and **U-X**) have much more HMGB1 than the wounds treated with saline (**A-D**, and **M-P**). KO-PRP: PRP from PLT HMGB1-KO mice; and GFP-PRP: PRP from GFP mice. White bars: 200 μm, Yellow bars: 50 μm (immunostaining).

### PRP from PLT-HMGB1-KO mice increases inflammation in wounded tendon

Collected patellar tendon tissues were analyzed for CD68, a marker of M1 pro-inflammatory macrophages (19, 20), using immunostaining. Overall, the levels of CD68 in the wound areas of PLT HMGB1-KO mice (**Fig. 5A-F**) were much higher than that of GFP mice (**Fig. 5G-L**). Many CD68 positive cells were found in wounded tendons of both groups of mice treated with saline (**Fig. 5A, B, 5G, H**) suggesting a high level of tissue inflammation. In PLT HMGB1-KO mice, GFP-PRP treatment decreased CD68 expression (**Fig. 5E, F**) compared to PRP from KO mice (**Fig. 5C, D**). However, in GFP mice, treatment with GFP-PRP significantly decreased positively stained CD68 cells (**Fig. 5K, L**) compared to the same group treated with PRP from KO mice (**Fig. 5I, J**). Taken together, these results suggest that ablation of PLT HMGB1 in PRP results in significant levels of CD68^+^ M1 macrophages in treated tendon tissues, while M1 cells are greatly reduced in tendons treated with normal PRP treatments. Semi-quantification supports these results (**Fig. 5M**), showing that GFP-PRP is able to reduce the level of CD68^+^ M1 macrophages in both transgenic mouse lines, while both saline and KO-PRP are largely similar in the level of CD68^+^ cells.

**Fig. 5.**
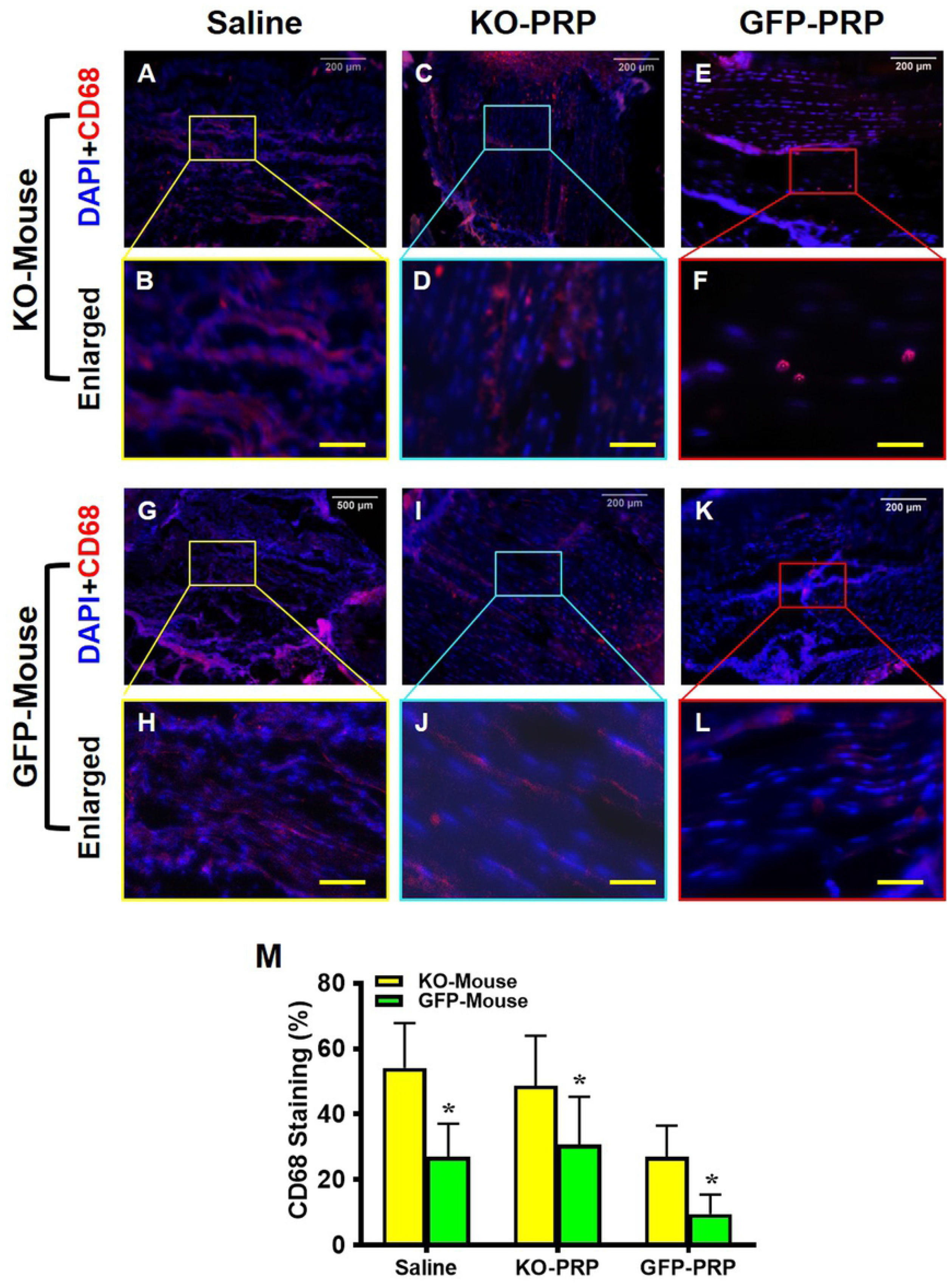
Macrophage marker CD68 expression is much higher in KO-PRP treated wound than GFP-PRP treated wound. Higher levels of CD68 are found in the wound areas of PLT HMGB1-KO mouse patellar tendons (**A-F**) than that of GFP mouse tendons (**G-L**). GFP-PRP treated wound areas (**E, F**, and **K, L**) have decreased CD68 expression compared with saline treated wounds (**A, B**, and **G, H**). Semi-quantification (**M**) confirms the results. *P < 0.01 (KO-PRP vs. GFP-PRP). KO-PRP: PRP from PLT HMGB1-KO mice; and GFP-PRP: PRP from GFP mice. White bars: 200 μm; Yellow bars: 50 μm (immunostaining).

### Stem cell marker expression is reduced in wounded mouse tendons treated with PRP from PLT-HMGB1-KO

HMGB1 have been shown to enhance tissue repair by recruiting resident stem cells (21). Thus, to assess the effect of PLT HMGB1 on resident stem cells patellar tendon tissues were immunostained for CD146 and CD73 stem cell marker expression (22, 23). GFP-PRP treatment of both transgenic mouse lines recruited stem cells to the wound areas, as shown by elevated CD146^+^ cells within GFP-PRP treated tendons (**Fig. 6E, F, 6K, L**) compared to tendons treated with KO-PRP (**Fig. 6C, D, 6I, J**). However, few CD146^+^ cells can be seen in the saline-treated tendons of PLT-HMGB1-KO mice (**Fig. 6A, B**) compared to saline treatment of GFP mice (**Fig. 6G, H**). Overall, higher levels of CD146^+^ cells were found in the tendons of GFP mice (**Fig. 6G-L**) compared to PLT HMGB1-KO tendons (**Fig. 6A-F**). Semi-quantification supports these results (**Fig. 6M**), showing that GFP-PRP treatment elevated CD146^+^ cells in both transgenic lines, with the highest levels found within GFP mice.

**Fig. 6.**
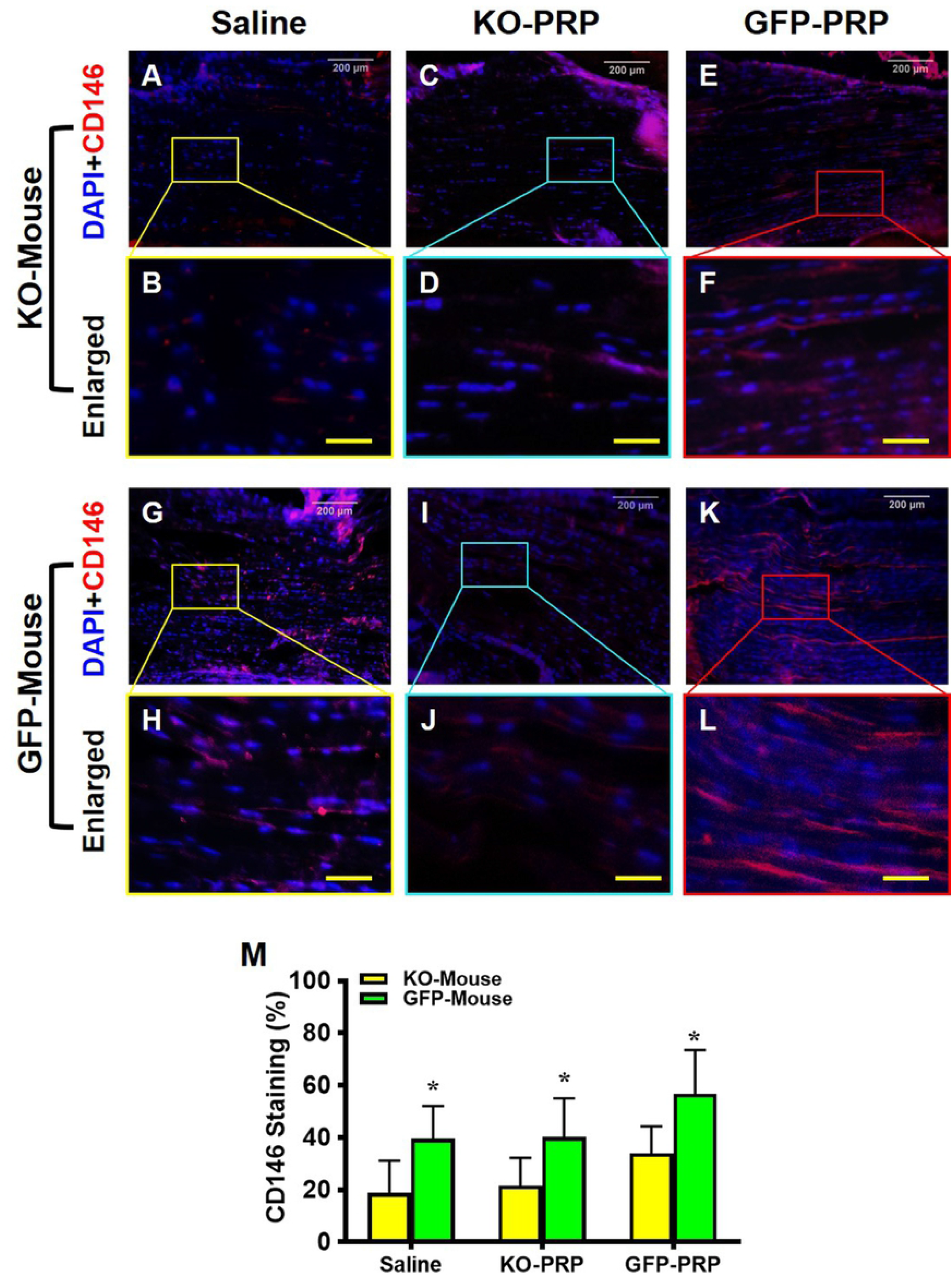
Stem cell marker CD146 expression is much lower in KO-PRP treated wound than GFP-PRP treated wound. Very low levels of CD146 are found in the wound areas of KO-PRP treated mice (**A-F**) compared GFP tendons which have much higher level of CD146 expression (**G-L**). GFP-PRP treated wound areas (**E, F and K, L**) have much more CD146 than the wounds treated with saline (**A-D and G-J**). Semi-quantification (**M**) confirms the results. *P < 0.01 (KO-PRP vs. GFP-PRP). KO-PRP: PRP from PLT HMGB1-KO mice; and GFP-PRP: PRP from GFP mice. White bars: 200 μm; Yellow bars: 50 μm (immunostaining).

Similar results were found with CD73 levels in wounded tendons (**Fig. 7**). Few CD73^+^ cells can be seen within saline-treated tendons of PLT HMGB1-KO mice (**Fig. 7A, B**), compared to saline treated GFP mice (**Fig. 7G, H**). Overall, both PLT HMGB1-KO PRP and GFP-PRP were able to increase the level of CD73^+^ cells but to different degrees. GFP-PRP treatment increased CD73^+^ cells in both mouse lines (**Fig. 7E, F, 7K, L**), surpassing the effect of PLT HMGB1-KO PRP on CD73 levels (**Fig. 7C, D, 7I, J**). Semi-quantification of CD73 staining supports these results (**Fig. 7M**), with GFP-PRP treatment producing elevated CD73^+^ cells in both transgenic lines with the highest levels in the treated tendons of GFP mice. Taken together, these results suggest that HMGB1 ablation in PLTs can negatively affect tendon wound healing by decreasing stem cell recruitment.

**Fig. 7.**
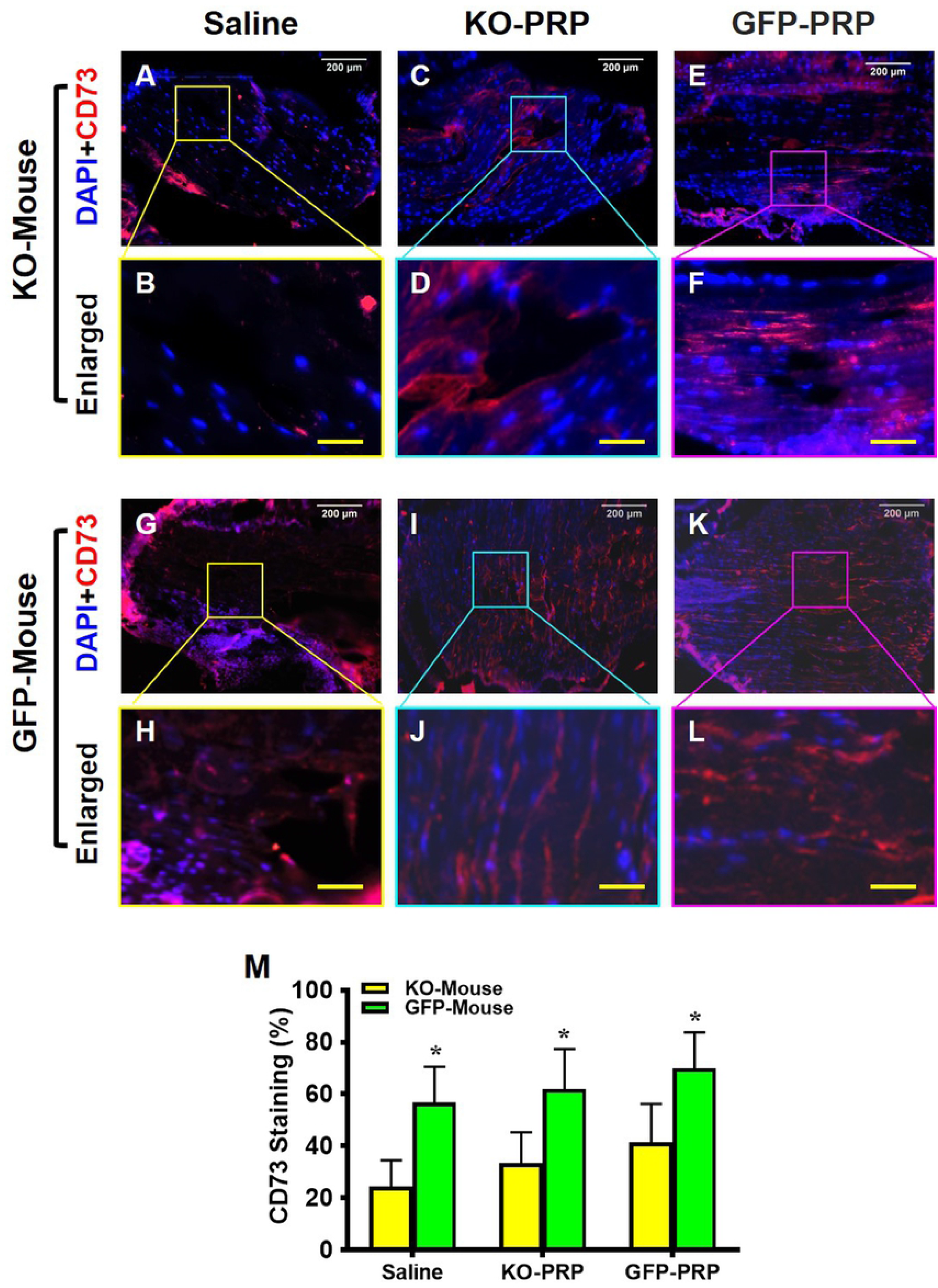
Stem cell marker CD73 expression is much lower in KO-PRP treated wound compared to GFP-PRP treated wound. Very low levels of CD73 are found in the wound areas of KO-PRP mouse patellar tendons (**A-F**) compared to GFP tendons, which have much higher levels of CD73 expression(**G-L**). However, both KO-PRP treated and GFP-PRP treated wound areas (**C-F**, and **I-L**) have more CD73 than the wounds treated with saline (**A, B**, and **G, H**). Semi-quantification (**M**) confirms the results. *P < 0.01 (KO-PRP vs. GFP-PRP). KO-PRP: PRP from PLT HMGB1-KO mice; GFP-PRP: PRP from GFP mice. White bars: 200 μm; Yellow bars: 50 μm (immunostaining).

### PRP from PLT-HMGB1-KO mice impairs collagen III production in wounded tendons

Collagen type III (Col III) has an important role in the healing process of tendon (24). Immunostaining for Col III was carried out to assess healing within wounded and treated patellar tendons. Overall, Col III expression was higher in the treated tendons of GFP mice (**Fig. 8G-L**) compared to tendons in PLT HMGB1-KO mice (**Fig. 8A-F**). Also, GFP-PRP treatment increased Col III levels in wounded tendons in both mouse lines (**Fig. 8E, F, 8K, L**), compared to PLT HMGB1-KO PRP treatment (**Fig. 8C, D, 8I, J**). The results indicate that in contrast to normal PLTs in GFP mice, HMGB1-ablated PLTs impair tendon wound healing due to decreased Col III levels. Semi-quantification supports this conclusion (**Fig. 8M**), with the GFP-PRP treatment surpassing the KO-PRP treatment in Col III production. Finally, similar to other results above, GFP-PRP treatment was able to produce the highest Col III production in GFP mice.

**Fig. 8.**
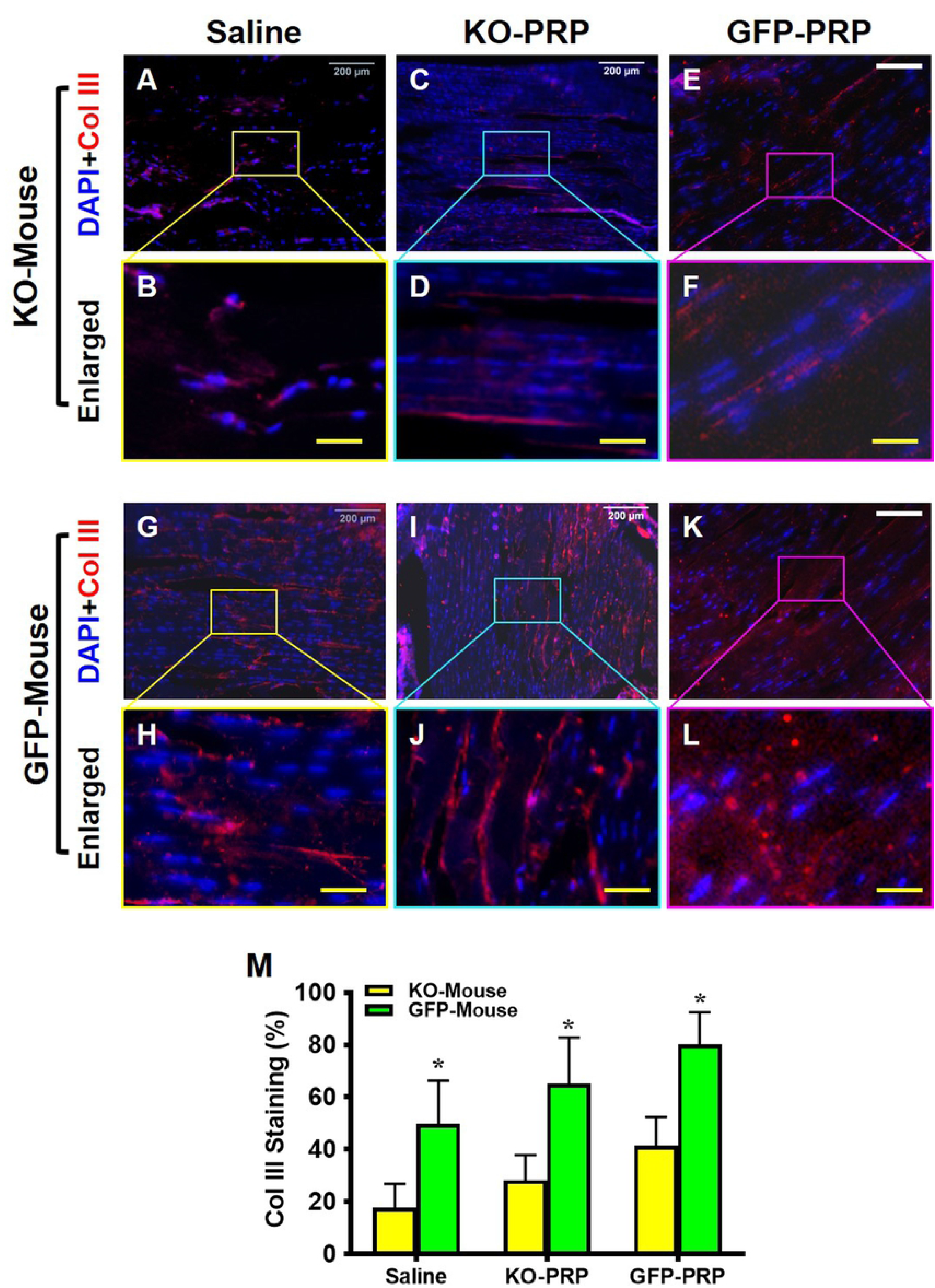
Tendon matrix marker collagen III expression is much lower in KO-PRP treated wound than GFP-PRP treated wound. Very low levels of collagen III are found in the wound areas of KO-PRP mouse patellar tendons (**A-F**) compared to GFP tendons with higher levels of Col III (**G-L**). KO-PRP and GFP-PRP treated wound areas (**C-F**, and **I-L**) have more Col III than the wounds treated with saline (**A, B**, and **G, H**). Semi-quantification (**M**) confirms the results. *P< 0.01 when KO-PRP treatment is compared to GFP-PRP treatment. KO-PRP: PRP from PLT HMGB1-KO mice; GFP-PRP: PRP from GFP mice. Col III: Collagen type III. White bars: 200 μm; Yellow bars: 50 μm (immunostaining).

## Discussion

This study investigated the effect of platelet-derived HMGB1 in PRP on wounded tendon healing using a transgenic mouse line with a specific platelet HMGB1 ablation. Our results have demonstrated that in an acute patellar tendon injury model treated with PRP from mice lacking platelet-derived HMGB1, tendon healing was impaired. Impaired healing was characterized by a decrease in local HMGB1 release from injured tissues, an increase in CD68^+^ M1 macrophage cells, a decrease in CD146^+^ and CD73^+^ stem cells, and a decrease in Col III content. In contrast, treatment with PRP from GFP-mice increased local HMGB1 concentrations in the wound area, reduced the recruitment of inflammatory cells to wounded tendons, increased the recruitment of stem cells, and increased Col III levels. Although the wounds treated with PLT HMGB1-KO-PRP healed faster than the wounds treated with saline, they healed much slower in comparison to wounds treated with GFP-PRP. This slowed healing is likely due to the ablation of PLT HMGB1 in these mice, with our results showing the expression of minimal amounts of PLT HMGB1 (**Fig. 1**) that may still facilitate some level of healing compared to the saline treatment. Our results also demonstrated that GFP-PRP treatment enhanced wounded tendon healing in both mouse lines, however to a greater extent in GFP mice than in PLT HMGB1-KO mice. In GFP mice, endogenous HMGB1 in addition to PLT HMGB1 within GFP-PRP were collectively involved in enhancing tendon healing. Thus, our results showed that HMGB1 in platelets is a critical factor in PRP treatment, and the efficacy of PRP treatment for tendon injuries in clinics may depend on the level of PLT HMGB1 within PRP preparations.

Tendon injury is one of the most common musculoskeletal injuries that can persist for years with poorly repaired tendon leading to further re-injury. In current clinical practices, PRP is widely used to treat tendon injuries (2, 25), with a number of studies showing that PRP treatment promotes the healing and physical function of wounded tendon (1, 2, 5, 26). These beneficial effects are attributed to the platelets in PRP, the role of which is well-established in wound healing and tissue repair (27). In fact, platelets are the “first responders” during wounding that trigger platelet activation and aggregation (28). Once activated, platelets release many factors (e.g. cytokine and growth factors) to enhance healing of injured tissues (29).

In particular, HMGB1, which is abundant in platelets (13, 15, 30), is also released by activated platelets. It has been shown that HMGB1 released by injured tissues promotes tissue repair by inducing migration and proliferation of stem cells (31-33). Locally released HMGB1 recruits bone marrow-derived mesenchymal stem cells (MSCs), and promotes the proliferation and differentiation of tissue-associated resident stem cells (11, 33, 34). For instance, vascularization of regenerating tissue is compromised in the absence of leukocyte HMGB1 in a mouse model lacking HMGB1 in the hematopoietic system, but vessel number and vascular area were significantly higher in the presence of leucocyte HMGB1 (35). This study indicated that leukocyte HMGB1 controls the nutrient and oxygen supply to the regenerating tissue. A study demonstrating the role of HMGB1 in muscle regeneration has shown that heterozygous HMGB1^+/–^ mice, which express 50% less HMGB1 when compared with wild type mice, have delayed muscle regeneration after acute injury (36). Fully reduced frHMGB1 has been gaining much attention as a chemoattractant that orchestrates tissue regeneration (18, 21). This subsequent increase in stem cells in response to HMGB1 administration suggested that the regenerative properties of HMGB1 were mediated by muscle stem cells and high expression of HMGB1 is required for optimal skeletal muscle regeneration (21). Exogenous administration of a single dose of HMGB1, either locally or systemically, promoted tissue repair by targeting endogenous stem cells. Using HMGB1^−/–^ mice, they identified the underlying mechanism as the transition of quiescent stem cells from G_0_ to G_Alert_, and these primed cells rapidly respond to appropriate activating factors released upon injury. These studies support our findings by highlighting the role of HMGB1 as a chemoattractant that promotes regeneration of wounded tendons by recruiting stem cells to injury site.

Macrophages are known to play an essential role in orchestrating inflammation and tissue repair (37). Macrophages secrete various growth factors and signaling molecules and are thus involved in the regulation of inflammation, wound healing, and tissue repair (38). Our findings show that active inflammation is present within wounds treated with PRP from PLT HMGB1-KO mice as indicated by the presence of CD68^+^ M1 macrophages (39), in contrast those treated with PRP from GFP mice. Inflammation can have a detrimental effect on healing. Chronic wounds fail to heal because they are stalled in an early inflammatory state during wound healing (40). For acute injuries, prolonged inflammation can lead to slow healing which may cause the wound to enter a chronic state and fail to heal (41). Thus, controlling inflammation may be an important step in preventing certain acute injuries from progressing to a chronic state. Research has shown that HMGB1 is able to mediate macrophage polarization (42). M1 macrophages are characterized by a proinflammatory phenotype which produces proinflammatory cytokines, phagocytosis of microbes, and initiate an immune response (19). M2 phenotype macrophages are a tissue-healing phenotype that releases more HMGB1, which may activate stem cells and promote tissue healing (43, 44). The switch from a proinflammatory M1 to a tissue-healing M2 phenotype in macrophages is an essential step in muscle regeneration to limit the inflammatory response (20, 45). Our results have shown that local HMGB1 is released in high levels in the wounded tendon matrix when treated with PRP from GFP mice as opposed to low levels of local HMGB1 release due to treatment with KO-PRP. This elevated level of local HMGB1 expression may help switch M1 macrophages to tissue healing M2 phenotype that may enhance healing (46). Further research however is needed to evaluate the effect of PLT HMGB1 on the M2 phenotype and on macrophage polarization in an acute tendon injury model.

As described above, many studies have demonstrated that HMGB1 activates endogenous stem cells to accelerate tissue regeneration (18, 21). Our results indicated that tendon wounds treated with PRP from mice lacking platelet-derived HMGB1 harbor a reduced number of CD146^+^ and CD73^+^ stem cells (**Fig. 6, 7**). Our results indicate that both platelet and local HMGB1 facilitates stem cell migration in normal wound healing. Our results also demonstrate high level of Col III expression in wounds treated by normal PRP suggesting normal wound healing is occurring, in contrast to the low level in KO mice PRP treated ones. Although Col III is not a major component of the normal tendon, it is believed to play an important role during the healing process because of its ability to form rapid crosslinks and stabilize the repair site (24).

Certain limitations are present within our study. Primarily, our resource for PLT HMGB1-KO mice is limited and as such our study only evaluated healing and repair at a single, relatively short timepoint (or 7 days post-injury) and only focused on the assessment of a limited number of cell markers. Further research is needed within a longer healing timeframe to further assess the effect of the ablation of PLT HMGB1 on tendon healing and repair.

In conclusion, this study has demonstrated that PLT HMGB1 within PRP plays an important role in healing wounded tendon by decreasing inflammation, increasing local HMGB1 levels, and recruiting stem cells to the wound area. These results provide the first evidence for the role of HMGB1 within PRP as a therapeutic treatment to promote tendon wound healing. Our findings suggest that the efficacy of PRP treatment for tendon injuries in clinics may depend on PLT HMGB1 within PRP preparations.

## Acknowledgements

We thank Dr. Bhavani P Thampatty for assistance in the preparation of this manuscript.

